# JNK pathway restricts DENV, ZIKV and CHIKV infection by activating complement and apoptosis in mosquito salivary glands

**DOI:** 10.1101/2020.03.01.972026

**Authors:** Avisha Chowdhury, Cassandra M. Modahl, Siok Thing Tan, Benjamin Wong Wei Xiang, Dorothée Missé, Thomas Vial, R. Manjunatha Kini, Julien Francis Pompon

## Abstract

Arbovirus infection of *Aedes aegypti* salivary glands (SGs) determines transmission. However, there is a dearth of knowledge on SG immunity. Here, we characterized SG immune response to dengue, Zika and chikungunya viruses using high-throughput transcriptomics. The three viruses regulate components of Toll, IMD and JNK pathways. However, silencing of Toll and IMD components showed variable effects on SG infection by each virus. In contrast, regulation of JNK pathway produced consistent responses. Virus infection increased with depletion of component Kayak and decreased with depletion of negative regulator Puckered. Virus-induced JNK pathway regulates complement and apoptosis in SGs via TEP20 and Dronc, respectively. Individual and co-silencing of these genes demonstrate their antiviral effects and that both may function together. Co-silencing either *TEP20* or *Dronc* with *Puckered* annihilates JNK pathway antiviral effect. We identified and characterized the broad antiviral function of JNK pathway in SGs, expanding the immune arsenal that blocks arbovirus transmission.

## Introduction

In recent decades, dengue (DENV), Zika (ZIKV) and chikungunya (CHIKV) viruses have emerged as global public health issues, with over 50% of the world population at risk for infection (Kraemer et al., 2019). DENV and ZIKV belong to the *Flavivirus* genus (Flaviviridae family), while CHIKV belongs to the *Alphavirus* genus (Togaviridae family). DENV infects an estimated 390 million people yearly, causing a wide range of clinical manifestations from mild fever to shock syndrome and fatal hemorrhage (Bhatt et al., 2013). ZIKV recently emerged as an epidemic virus, infecting 1.5 million people over the past five years (Hamel et al., 2016). Although ZIKV infection is mostly asymptomatic, it can result in life-debilitating neurological disorders including Guillain-Barré syndrome in adults and microcephaly in prenatally-infected newborns. CHIKV emerged as an epidemic virus in 2004, and has infected more than 6 million people (Silva & Dermody, 2017). It causes mild fever but can result in musculoskeletal inflammation, leading to long-term polyarthralgia. Amplified by urban growth, climate change and global travel, arboviral outbreaks are unlikely to recede in the near future (Mayer et al., 2017).

DENV, ZIKV and CHIKV are primarily transmitted by *Aedes aegypti* mosquitoes. In the absence of efficient vaccines (Wilder-Smith & Gubler, 2015) and curative drugs(Barrows et al., 2018), targeting this common vector is the best available strategy to control the spread of all three viruses. However, current vector control methods that rely on chemical insecticides are not effective in preventing outbreaks (Wilder-Smith et al., 2017), partly due to insecticide resistance (Liu, 2015). A novel strategy employing *Wolbachia* to reduce virus transmission by mosquitoes has been deployed as a trial in several countries (Anders et al., 2018). However, its long-term efficacy may be compromised by converging bacteria and virus evolution (Ritchie et al., 2018). Other promising approaches utilize genetic engineering technology to develop refractory vector populations (Kistler et al., 2015). Mosquito innate immunity can drastically reduce virus transmission, and characterization of mosquito immune pathways and mechanisms will aid in identifying gene candidates for transformation.

Immunity in midguts, the first organ to be infected following a blood meal, has been extensively studied by using transcriptomics. DENV and ZIKV activate the Toll, Immune Deficiency (IMD) and Janus Kinase (JAK)/Signal Transduction and Activators of Transcription (STAT) immune pathways (Angleró-Rodríguez et al., 2017; Sim et al., 2013; Souza-Neto et al., 2009; Xi et al., 2008). Selective gene silencing studies have demonstrated the anti-DENV impact of Toll and JAK/STAT pathways. However, JAK/STAT transgenic activation was not effective in reducing ZIKV and CHIKV infection (Jupatanakul et al., 2017). Immune effectors downstream of these pathways include antimicrobial peptides (AMPs) and thioester containing proteins (TEPs) (Angleró-Rodríguez et al., 2017; Jupatanakul et al., 2017; Ramirez & Dimopoulos, 2010; Sim et al., 2013; Souza-Neto et al., 2009). AMPs can have direct antiviral activity (Luplertlop et al., 2011), while TEPs that belong to the complement system tag pathogens for lysis, phagocytosis or melanization (Blandin, 2004; Xiao et al., 2014). Additionally, the RNA interference (RNAi) pathway cleaves viral RNA genomes (Sanchez-Vargas et al., 2004). However, its impact against arbovirus infection is uncertain (Olmo et al., 2018). In *Drosophila melanogaster,* the Jun-N-terminal Kinase (JNK) pathway regulates a range of biological functions including immunity and apoptosis (Kockel et al., 2001). In *Anopheles gambiae,* the JNK pathway mediates an anti-malaria response through complement activation (Garver et al., 2013). Currently, the impact of JNK pathway on arbovirus infection remains unexplored.

Following midgut invasion, arboviruses propagate to remaining tissues, including salivary glands (SGs), from where they are expectorated with saliva during subsequent bites. Despite the critical role of SGs in transmission, only three studies have examined DENV-responsive differential gene expression in SGs. These studies revealed activation of the Toll and IMD pathways (Bonizzoni et al., 2012; Luplertlop et al., 2011; Sim et al., 2012) and identified the anti-DENV functions of Cecropin (Luplertlop et al., 2011), putative Cystatin and ankyrin-repeat proteins (Sim et al., 2012). Here, we characterized the SG immune response to DENV, ZIKV and CHIKV. We performed the first high throughput RNA-sequencing (RNA-seq) in infected SGs and observed differentially expressed genes (DEGs) related to immunity, apoptosis, blood-feeding and lipid metabolism. Using gene silencing, we discovered that upregulated components of the Toll and IMD pathways had variable effects against DENV, ZIKV and CHIKV. However, silencing of a JNK pathway upregulated component increased, and silencing of a negative regulator decreased infection by the three viruses in SGs. Further, we show that the JNK pathway is activated by all viruses and triggers a cooperative complement and apoptosis response in SGs. This work identifies and characterizes the JNK antiviral response that reduces DENV, ZIKV and CHIKV in *A. aegypti* SGs.

## Results

### Transcriptome regulation by DENV, ZIKV and CHIKV in SGs

SGs were collected at 14 days post oral infection (dpi) with DENV and ZIKV, and at seven dpi with CHIKV (Supplementary Figure 1) to account for variability between virus extrinsic incubation periods (EIP) (Mbaika et al., 2016; Salazar et al., 2007). Differentially expressed genes (DEGs) were calculated with DESeq2, edgeR and Cuffdiff 2, and showed little overlap among the algorithms (Supplementary Figure 2). To validate DEGs and select which software to use, we quantified the expression of 10 genes in a biological repeat with RT-qPCR and compared these values to the output from each algorithm. RNA-seq gene expression obtained with DESeq2 correlated best with RT-qPCR values (DENV: r^2^ = 0.69; ZIKV: r² = 0.81; CHIKV: r² = 0.79; Supplementary Figure 3), and only these DEGs are discussed.

The SG transcriptome was the most regulated by CHIKV infection (966 DEGs), followed by ZIKV (396) and DENV (202) (Figure 1a; Supplementary Table 1). Only 19 DEGs were common amongst the three virus infections (Supplementary text), indicating a virus-specific transcriptome response. Comparison between DEGs from the current DENV infection and the three previous DENV studies with SGs collected at 14 dpi showed little overlap among them (Supplementary Figure 4) (Bonizzoni et al., 2012; Luplertlop et al., 2011; Sim et al., 2012). However, despite technical variations (e.g. mosquito colony, virus strain, transcriptomics technology), DEGs belonged to similar functional groups across the studies. We observed that immunity, apoptosis, blood-feeding and lipid metabolism related genes were highly regulated by DENV, ZIKV and CHIKV (Figure 1b; Supplementary text).

**Figure 1.**
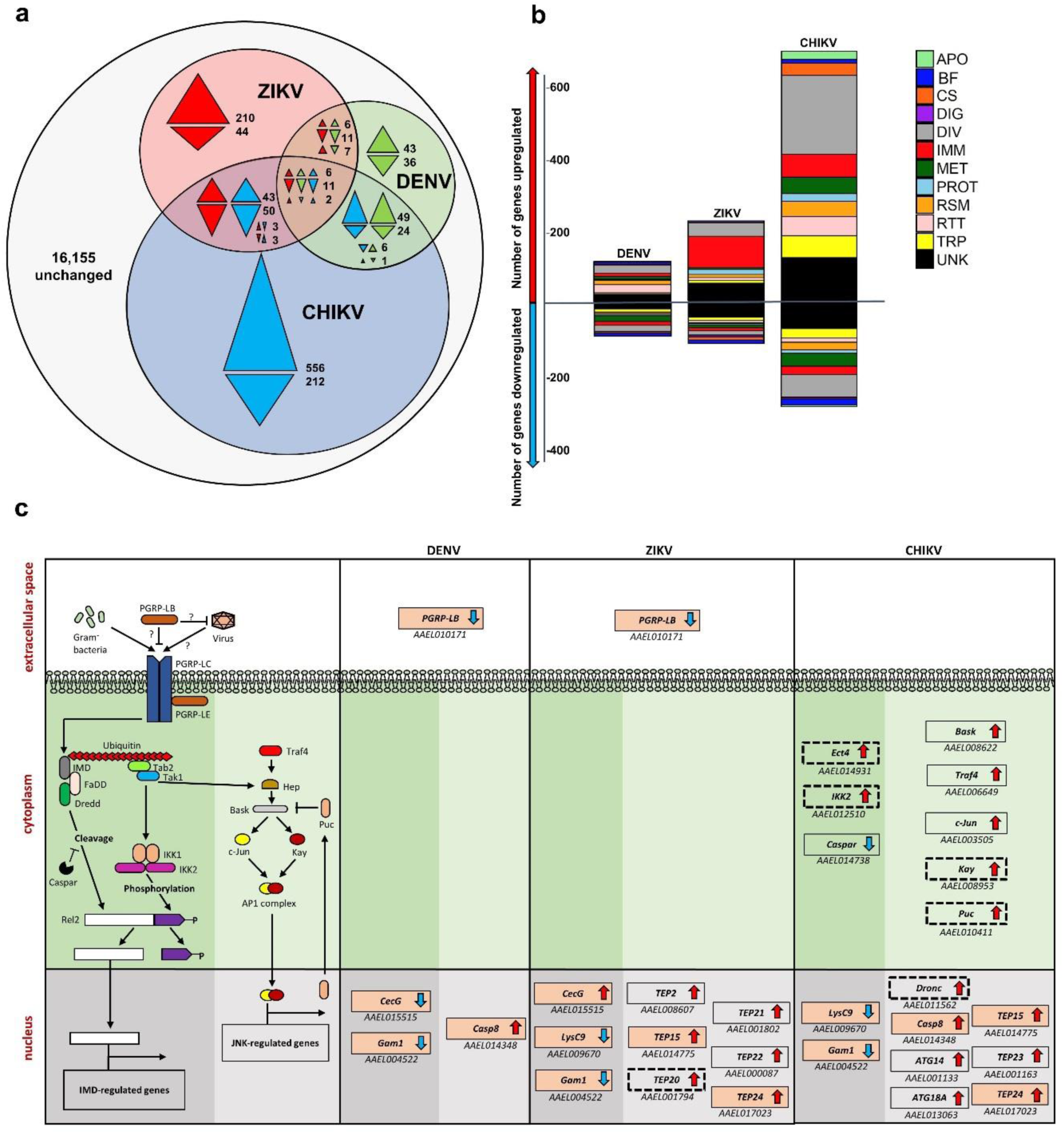
DENV, ZIKV and CHIKV transcriptome regulation in *A. aegypti* salivary glands. **a**, Venn diagram shows numbers of uniquely and commonly regulated differentially expressed genes (DEGs) in DENV (green), ZIKV (red) and CHIKV (blue) infected salivary glands. Colorless area shows total number of unchanged genes. Arrows indicate the direction of regulation for the corresponding color code. **b**, Functional annotation of DEGs in DENV, ZIKV or CHIKV infected salivary glands. APO, apoptosis; BF, blood feeding; CS, cytoskeleton and structure; DIG, digestion; DIV, diverse functions; IMM, immunity; MET, metabolism; PROT, proteolysis; RSM, redox, stress and mitochondria; RTT, replication, transcription and translation; TRP, transport; UNK, unknown functions. **c,** Transcriptomic regulation of the IMD and JNK pathways in DENV, ZIKV and CHIKV infected salivary glands. First column represents a scheme of the JNK and IMD pathways partially, modified from (Sim et al., 2014). In the following columns, boxes indicate DEGs with AAEL number below for DENV, ZIKV and CHIKV. Arrows indicate the direction of regulation. Pink-filled boxes indicate genes regulated by more than one virus. Boxes with dotted line indicate DEGs selected for functional studies. Dark and light shaded green areas differentiate IMD from JNK pathways.

A high proportion of DEGs were related to immunity with DENV, ZIKV and CHIKV modulating 20, 118 and 86 immune genes, respectively (Figure 1b; Supplementary Table 1). Differential regulation of the RNAi, Toll, IMD, JAK/STAT and JNK pathways were observed. The RNAi pathway is initiated when Dicer2 (Dcr2) recognizes and cleaves viral-derived dsRNA into small-interfering RNAs (siRNA) (Supplementary Figure 5). siRNAs are loaded onto the RNA-induced silencing complex (RISC) that includes Argonaute2 (Ago2). RISC unwinds siRNA to allow binding to complementary viral RNA fragments and cleavage (Sanchez-Vargas et al., 2004). Strikingly, *Dcr2* was upregulated by all viruses, while *Ago2* was upregulated by DENV (Supplementary Figure 5), suggesting activation of RNAi. The Toll pathway is triggered when extracellular pattern recognition receptors (PRR) (e.g. PGRP and GNBP) bind to pathogen-derived ligands and activate a proteolytic cascade that leads to activation of pro-Spätzle to Spätzle by Spätzle processing enzyme (SPE) (Supplementary Figure 6) (DeLotto & DeLotto, 1998). Spätzle binding to transmembrane receptor Toll induces a cytoplasmic cascade that leads to nuclear translocation of NF-κB transcription factor Rel1a to initiate effector gene transcriptions. Among PRRs, we observed that ZIKV and CHIKV upregulated *GNBPA1* and downregulated *GNBPB6*, while ZIKV also upregulated *PGRPS1* and CHIKV downregulated *GNBP2* (Supplementary Figure 6). Numerous serine proteases [e.g. CLIPs including Snake-likes (Snk-like) or Easter-likes (Est-like)] and serine protease inhibitors (Serpins) were upregulated by the viruses, except for *Snk-like* AAEL002273 and *CLIPB41* downregulation by CHIKV, and *SPE* and *CLIPB37* downregulation by DENV. *SPE* downregulation by DENV is reminiscent of downregulation of Toll pathway components by subgenomic flaviviral RNA (sfRNA) in SGs (Pompon et al., 2017). The IMD pathway, similarly to Toll, is triggered by microbial ligand recognition by PRRs, which activate PGRP-LC transmembrane protein (Figure 1c). In the ensuing cytoplasmic cascade, NF-κB transcription factor Rel2 is phosphorylated by IKK2 and cleaved by DREDD to induce its nuclear translocation. IMD pathway is repressed by Caspar (Sim et al., 2014), an Ectoderm-expressed 4-containing module (Ect4) (Akhouayri et al., 2011) and PGRP-LB (Gendrin et al., 2017). We observed an induction of IMD pathway by DENV and ZIKV through *PGRP-LB* downregulation, and by CHIKV through *Caspar* downregulation and *IKK2* upregulation (Figure 1c). *Ect4* upregulation by CHIKV may have controlled the pathway activation. The JAK/STAT pathway is triggered by binding of Unpaired (Upd) to transmembrane protein Domeless (Dome), which dimerizes and phosphorylates Hopscotch (Hop) (Supplementary Figure 7) (Sim et al., 2014). The activated Dome/Hop complex phosphorylates STATs, which upon dimerization translocate to nucleus and initiate transcription. The phosphorylation of STATs is controlled by Suppressor of Cytokine Signaling (SOCS) family of inhibitors (Callus & Mathey-Prevot, 2002). We observed an upregulation of *SOCS36E* by CHIKV (Supplementary Figure 7), suggesting inhibition of the pathway. The JNK pathway is triggered by Hemipterous (Hep) phosphorylation through tumor necrosis-associated factor 4 (Traf4) or Tak1, the latter being an IMD pathway component (Figure 1c). Hep activates Basket (Bask) that phosphorylates c-Jun and Kayak (Kay) transcription factors. Activated c-Jun and Kay dimerize to form the activator protein-1 (AP-1) complex, which translocates to nucleus and induces transcription. *Puckered* (*Puc*) transcription is induced by the JNK pathway and represses Bask (Martín-Blanco et al., 1998), acting as a feedback loop. Intriguingly, CHIKV upregulated most of the JNK pathway components (i.e. *Bask*, *c-Jun*, *Kay* and *Puc*; Figure 1c), indicating activation of the pathway in SGs. Furthermore, *Traf4* upregulation by CHIKV suggested an IMD-independent activation.

### Upregulated components from JNK but not from Toll and IMD pathway reduce DENV, ZIKV and CHIKV

To determine how immune response influences DENV, ZIKV and CHIKV multiplication in SGs, we silenced nine previously uncharacterized immune genes that were upregulated by at least one of the viruses (Table 1). They included four protease genes that initiate the Toll pathway (*CLIPB13A, CLIPB21,* one of the *Est-like,* one of the *Snk-like*), two genes from the cytoplasmic signaling of the IMD pathway (*IKK2* and *Ect4*), one transcription factor gene of the JNK pathway (*Kay*), and two putative immune genes - *Galectin-5* (*Gale5*) and a juvenile hormone induced gene (*JHI*). *Gale5* shows antiviral function against O’nyong-nyong in *A. gambiae* (Waldock et al., 2012), while juvenile hormone treatment regulates immune gene expression in *D. melanogaster* (Flatt et al., 2008) and *JHI* was upregulated by all three viruses in our study (Table 1). RNAi-mediated gene silencing efficacy ranged from 35-85 % in SGs (Supplementary Figure 8a) and did not affect mosquito survival (Supplementary Figure 9a). To bypass the midgut barrier, we intrathoracically inoculated mosquitoes with a non-saturating inoculum of DENV, ZIKV or CHIKV, resulting in 70-80% infected SGs (Supplementary Figure 10). This permitted the evaluation of an increase or decrease in infection upon gene silencing. At 10 days post inoculation, viral infection in SGs was measured using two parameters, infection rate and infection intensity. Infection rate was defined as the percentage of infected SGs out of 20 inoculated ones and represents dissemination of intrathoracically injected viruses into SGs. Infection intensity was calculated as viral genomic RNA (gRNA) copies per individual infected SGs and indicates virus replication. Of note, the two infection parameters do not reflect isolated biological phenomenon. For instance, a negative impact on infection intensity may lower infection rate due to virus clearance.

**Table 1.**
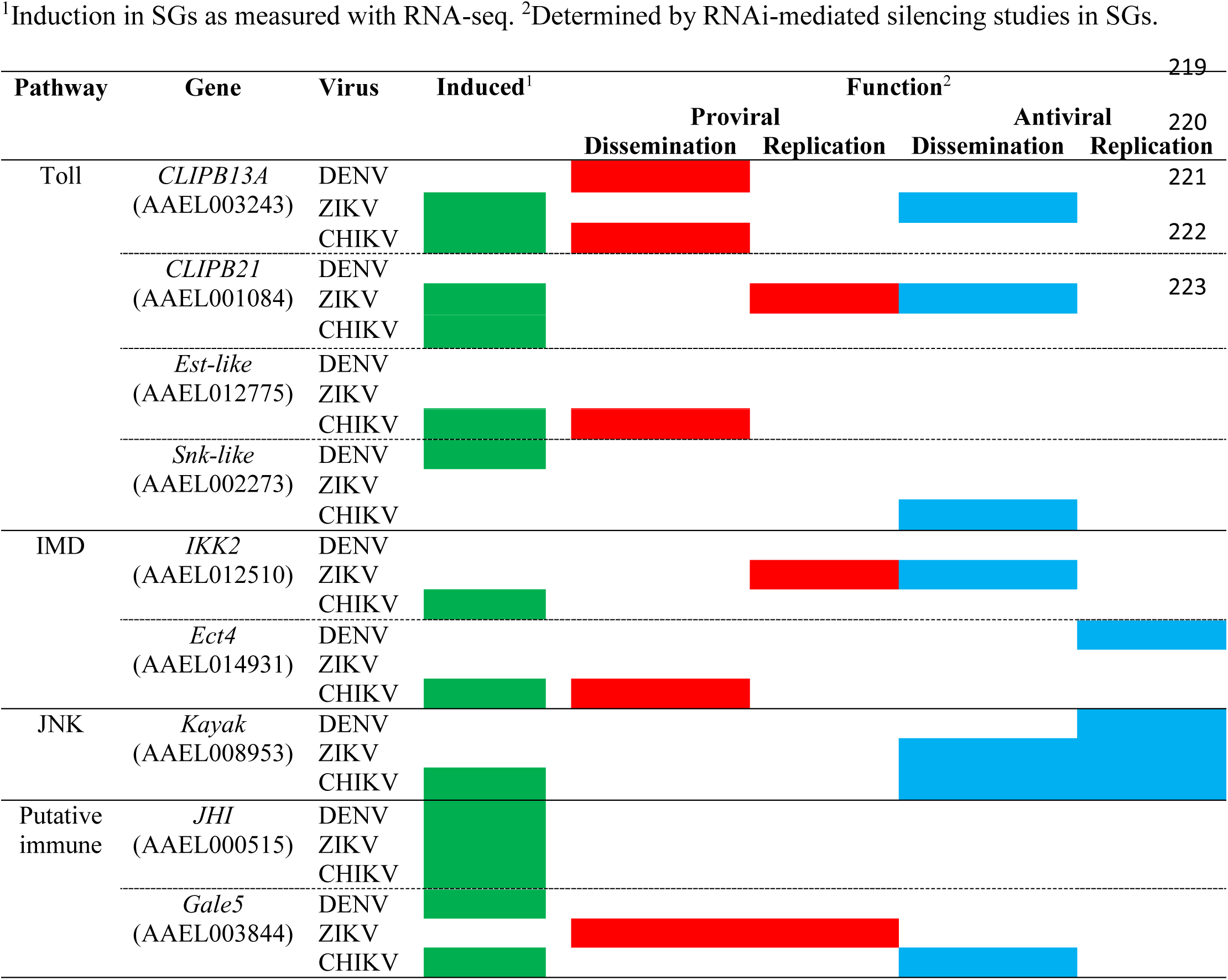
Impact of immune response on DENV, ZIKV and CHIKV infection in salivary glands

Silencing of most of the immune genes had a virus-specific effect on infection intensity and infection rate (Figure 2a-c and Table 1). Silencing of *CLIPB13A* increased ZIKV infection rate, but decreased DENV and CHIKV infection rates. This suggests that *CLIPB13A* hinders ZIKV dissemination and facilitates DENV and CHIKV dissemination into SGs. However, it has a minor role in regulating virus replication in SGs. *CLIPB21* silencing increased ZIKV infection rate but decreased its infection intensity. *Est-like* silencing decreased, whereas *Snk-like* silencing increased CHIKV infection rate. Overall, Toll pathway upregulated components showed mostly a virus-specific impact on dissemination into SGs (Figure 2a-c and Table 1). Silencing of *IKK2* increased infection rate and decreased infection intensity for ZIKV. *Ect4* silencing increased DENV infection intensity and decreased CHIKV infection rate. Overall, IMD pathway response appears both proviral and antiviral (Figure 2a-c and Table 1). *Gale5* silencing decreased both ZIKV infection intensity and infection rate, and enhanced CHIKV infection rate. Although *JHI* was upregulated by all viruses, it had no effect on any infections (Figure 2a-c and Table 1). The silencing studies reflect a complex interaction between transcriptomic response and antiviral functions (Table 1) with the notable exception of Kay. *Kay* silencing increased infection rate to 100% for both ZIKV and CHIKV, and increased infection intensities by 8.6-, 6.75- and 17.65-fold for DENV, ZIKV and CHIKV, respectively (Figure 2a-c and Table 1). These results reveal a broad antiviral function of the JNK pathway in SGs. To determine whether JNK pathway also restricts virus in midguts, we orally infected *Kay-*silenced mosquitoes with ZIKV (Supplementary Figure 8b). At seven dpi, while infection rate was already saturated at 100% in the control, infection intensity increased in midguts (Figure 2d). Overall, our functional studies discovered the antiviral and proviral roles of several immune-related genes and established that JNK pathway has a broad ubiquitous antiviral function.

**Figure 2.**
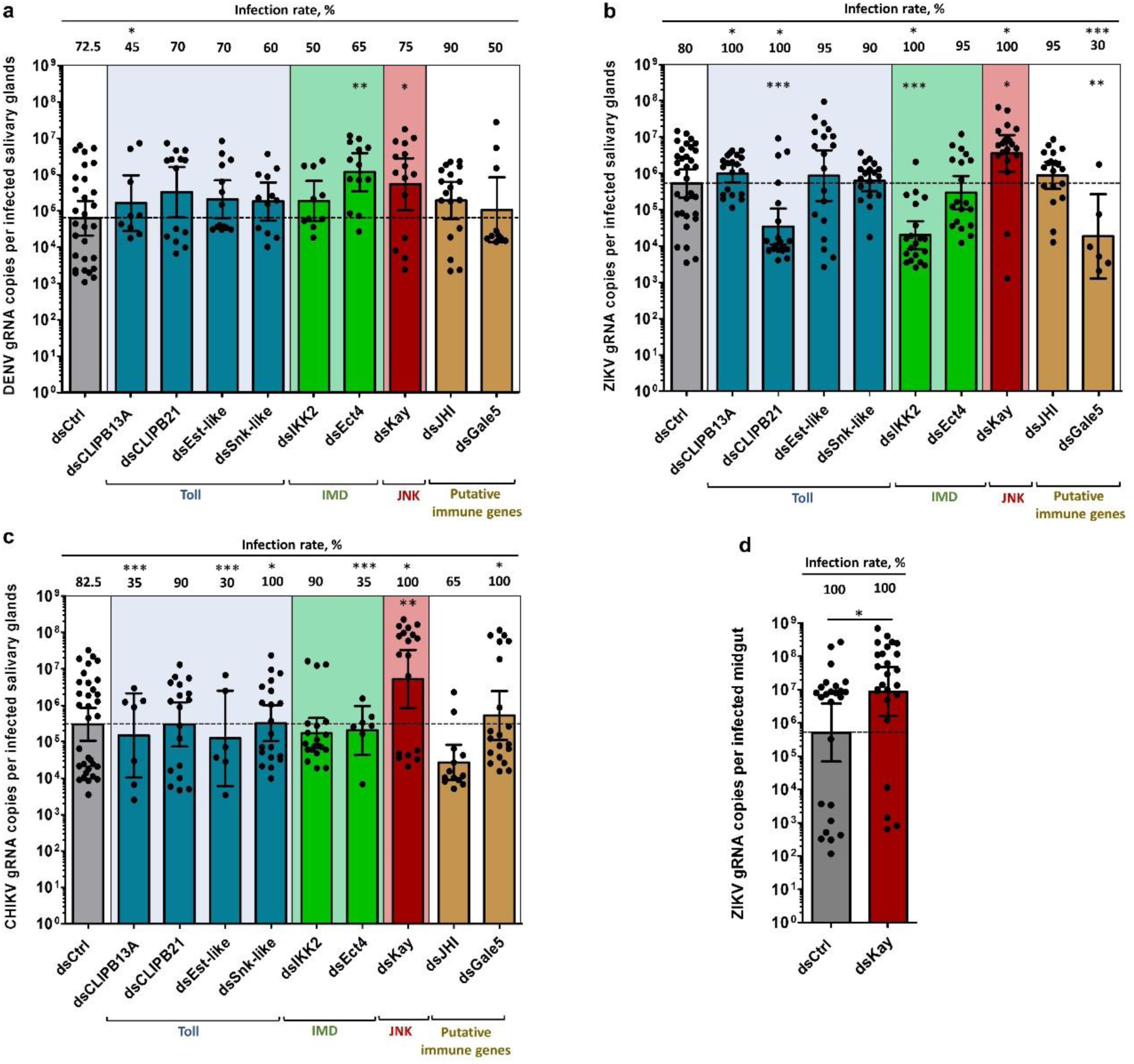
Kayak depletion but not Toll or IMD component depletion increases salivary glands infection by DENV, ZIKV and CHIKV, and midgut infection by ZIKV. Four days post dsRNA injection, mosquitoes were infected by intrathoracic inoculation with DENV, ZIKV or CHIKV, or by oral feeding on ZIKV. Viral genomic RNA (gRNA) was quantified at 10 days post inoculation in salivary glands and 7 days post oral infection in midguts. **a-c,** Effect of immune-related gene silencing on gRNA copies and infection rate in salivary glands infected with (**a)** DENV, (**b**) ZIKV, and (**c**) CHIKV. **d**, Impact of *Kayak* silencing on gRNA number and infection rate in ZIKV-infected midgut. Bars show geometric mean ± 95% C.I. from 20 individual pairs of salivary glands or 25 individual midguts. Each dot represents one sample. ds*Ctrl*, dsRNA against LacZ; ds*CLIPB13A*, dsRNA against CLIP domain serine protease B13A; ds*CLIPB21*, dsRNA against CLIP domain serine protease B21; ds*Est-like*, dsRNA against Easter-like; ds*Snk-like*, dsRNA against Snake-like; ds*IKK2*, dsRNA against Inhibitor of nuclear factor kappa-B kinase; ds*Kay*, dsRNA against Kayak; ds*Ect4*, dsRNA against Ectoderm expressed-4; ds*JHI*, dsRNA against Juvenile hormone inducible; ds*Gale5*, dsRNA against Galectin 5. *, p-value < 0.05; **, p-value < 0.01; ***, p-value < 0.001, determined by post hoc Dunnett’s with dsCtrl, unpaired t-test or Z-test for infection rate.

### JNK pathway is induced by DENV, ZIKV and CHIKV and reduces infection in SGs

SG transcriptomics showed that *Kay* was induced by CHIKV at seven dpi, but not by DENV or ZIKV at 14 dpi (Figure 1c). To test whether JNK pathway is activated by the three viruses, we monitored the kinetics of *Kay* expression in SGs after oral infection with either of the three viruses. *Kay* was similarly induced by DENV, ZIKV and CHIKV at three and seven dpi, but not at 14 dpi (Figure 3a), corroborating the RNA-seq data. We quantified *Puc* expression, which is induced as a negative regulator (Martín-Blanco et al., 1998). Although not regulated at three and 14 dpi, *Puc* expression was increased at seven dpi for all three viruses (Figure 3b). We then tested the impact of *Puc* silencing in ZIKV-inoculated mosquitoes (Supplementary Figure 8c). *Puc* silencing did not alter mosquito survival (Supplementary Figure 9b). Although SG infection rate was not affected, SG infection intensity was decreased by 8.85-fold at 10 days post inoculation (Figure 3c). Altogether our data demonstrate that the JNK pathway is induced by DENV, ZIKV and CHIKV at a time that corresponds to the onset of infection in SGs (Salazar et al., 2007) and further support the JNK antiviral function.

**Figure 3.**
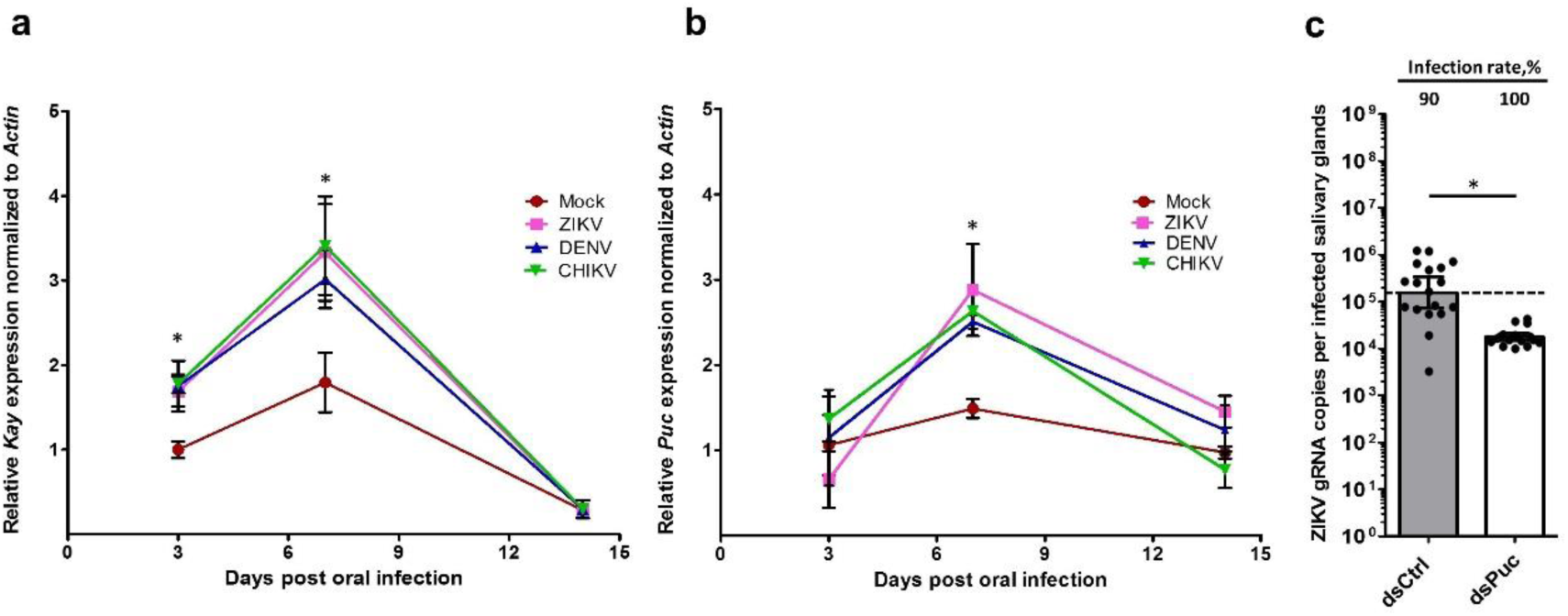
*Kayak* and *Puckered* expressions are induced by DENV, ZIKV and CHIKV infection, and Puckered depletion restricts ZIKV infection in salivary glands. **a,** *Kayak* (*Kay*) and **b,** *Puckered* (*Puc*) expressions in salivary glands at 3, 7 and 14 days post oral infection with DENV, ZIKV and CHIKV. Gene expression was quantified in pools of 10 salivary glands. *Actin* expression was used for normalization. Bars show arithmetic means ± s.e.m. from three biological replicates. **c**, Effect of Puc depletion on infection in salivary glands at 7 days post ZIKV inoculation. Four days prior infection, mosquitoes were injected with dsRNA. Bars show geometric means ± 95% C.I. from 20 individual salivary glands. Each dot represents one sample. *, p-value < 0.05, as determined by Dunnett’s test within time points with mock infection as control (a, b) or unpaired t-test (c).

### The JNK pathway mediates an antiviral complement and apoptosis response in SGs

JNK pathway can regulate complement system (Garver et al., 2013), apoptosis (McEwen, 2005) and autophagy (Wu et al., 2009). To determine whether JNK pathway induces these functions in SGs, we monitored the impact of *Kay* silencing on expressions of four TEPs (*TEP20*, *TEP24*, *TEP15* and *TEP2*), two pro-apoptotic (*Caspase8* and *Dronc*) and two autophagy-related (*ATG14* and *ATG18A*) genes at 10 dpi with ZIKV. All these genes were upregulated by infection in the transcriptomic data (Figure 1c; Supplementary Table 1). Control mosquitoes were injected with control dsRNA (dsCtrl). *Kay* silencing significantly reduced expressions of *TEP20* and *Dronc*, and moderately reduced *TEP15* and *TEP24* (Figure 4a). *TEP2* expression was increased and the two *ATGs* and *Caspase 8* were unaffected. These results suggest that upon infection the JNK pathway induces the complement system through TEPs, and apoptosis through Dronc, but not Caspase8. We then tested the antiviral function of TEP20 and Dronc in SGs by challenging either *TEP20-* or *Dronc-*silenced mosquitoes (Supplementary Figure 8d) by ZIKV intrathoracic inoculation. Similar to *Kay*-silencing, *TEP20-* and *Dronc*-silencing increased SG infection intensity by 21-fold to 2.6 x 10^6^ gRNA and by 12-fold to 1.3 x 10^6^ gRNA, respectively (Figure 4b). Since both complement and apoptosis can interact for cell clearance (Fishelson, 2001), we determined whether TEP20 and Dronc act in the same antiviral pathway by evaluating their synergistic effect when both were co-silenced (Supplementary Figure 8e). We did not observe a clear difference in SG infection between *TEP20* and *Dronc* individual or co-silencing (Figure 4c,d). As the infection conditions did not saturate the infection intensity (higher inoculum resulted in 10^8^ gRNA per infected SG; Supplementary Figure 10), the lack of synergism when TEP20 and Dronc are co-silenced suggests that TEP20 and Dronc function in the same antiviral pathway. To show that TEP20 and Dronc mediate the JNK antiviral response in SGs, we induced JNK pathway by silencing of *Puc* and co-silenced *TEP20* or *Dronc* before ZIKV inoculation. Control mosquitoes were injected with the same quantity of dsCtrl or dsRNA against *Puc*. While infection intensity decreased upon *Puc* silencing, co-silencing of *Puc* with *TEP20* or *Dronc* restored viral gRNA copies to the level in dsCtrl-injected SGs (Figure 4d). Altogether, these results demonstrate that the JNK pathway is induced by DENV, ZIKV or CHIKV in SGs and eliminates viruses through the complement system and apoptosis.

**Figure 4.**
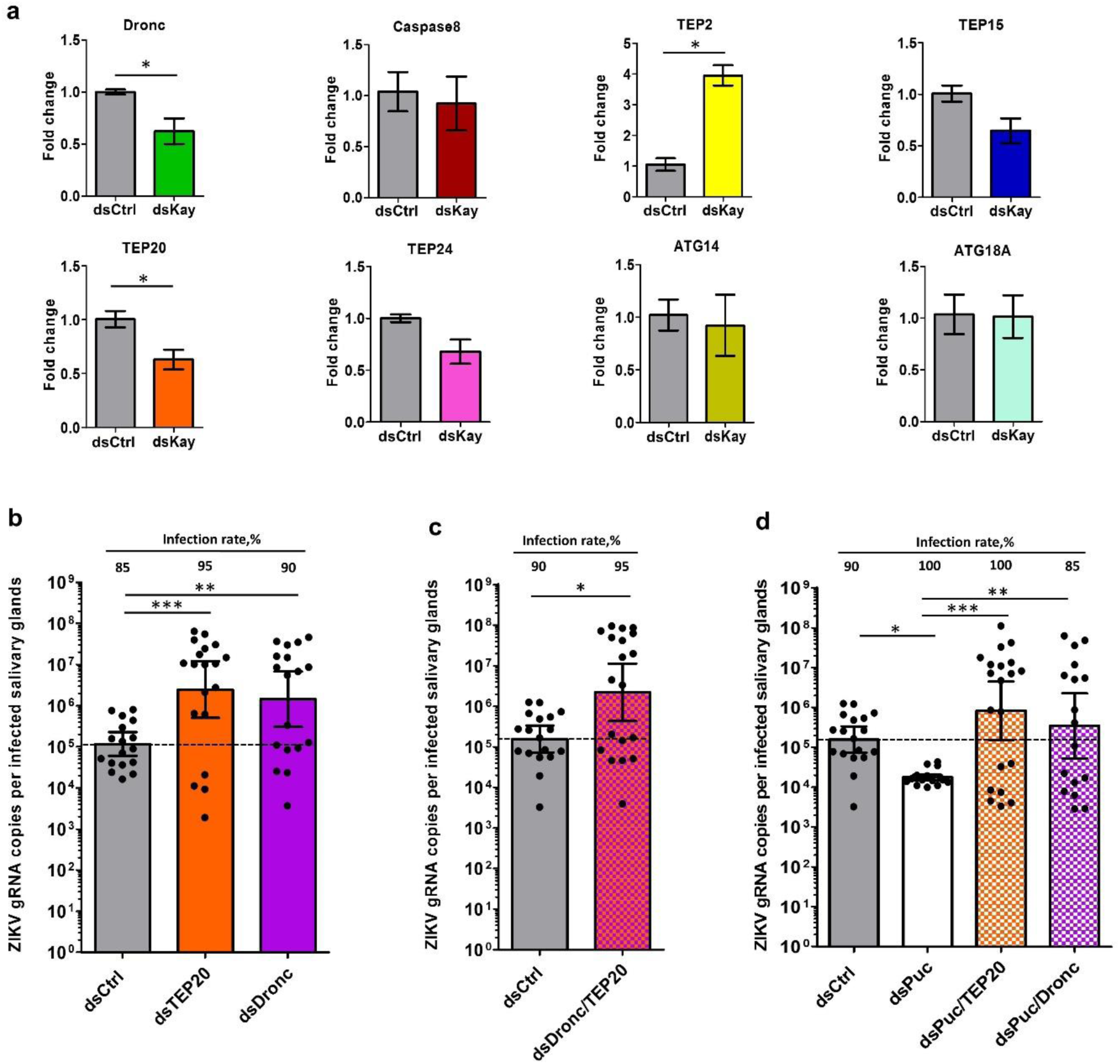
Virus-induced JNK pathway activates an antiviral response through complement and apoptosis. **a**, Impact of Kayak depletion on expression of genes related to complement (*TEP20*, *TEP24*, *TEP15* and *TEP2*), apoptosis (*Caspase8* and *Dronc*) and autophagy (*ATG14* and *ATG18A*) at 10 days post ZIKV oral infection. Gene expression was quantified in pools of 10 SGs. *Actin* expression was used for normalization. Bars show arithmetic means ± s.e.m. from three biological replicates. **b-d**, Effect of depletion of (**b**) Dronc or TEP20 alone, (**c**) Dronc and TEP20 simultaneously, and (**d**) Puc alone, or Puc and Dronc simultaneously, or Puc and TEP20 simultaneously on SG infection at 7 days post ZIKV inoculation. Four days prior infection, mosquitoes were injected with dsRNA. Bars show geometric means ± 95% C.I. from 20 individual salivary glands. Each dot represents one sample. *TEP,* Thioester-containing protein*; ATG,* Autophagy related gene; *Kay*, Kayak; *Puc*, Puckered; Ctrl, control. *, p-value < 0.05; **, p-value < 0.01; ***, p-value < 0.001, as determined by unpaired t-test (a) or Dunnett’s test with dsCtrl as control (b-d).

## Discussion

Dengue, Zika and chikungunya are widespread mosquito-borne diseases primarily transmitted by *A. aegypti*. Disease mitigation through insecticide-based vector control has mostly remained ineffective to prevent outbreaks (Wilder-Smith & Gubler, 2015). Currently, insecticide-free vector-control strategies are being extensively investigated. The improved understanding of the vector-virus interaction at the molecular level proposed here paves the way to manipulate mosquito biology to make them refractory towards arboviruses. This study discovers the antiviral function of the JNK pathway in SGs against three major arboviruses, DENV, ZIKV and CHIKV belonging to two families. Further, we determine that the antiviral response is mediated through a combined action of complement system and apoptosis.

Several studies have functionally characterized the immune response in *A. aegypti* midgut, the first barrier in the mosquito body, determining the antiviral function of Toll, IMD and JAKSTAT pathways (Angleró-Rodríguez et al., 2017; Bonizzoni et al., 2012; Dong et al., 2017; Sim et al., 2013). However, only a couple of studies tested the impact of SG (exit barrier) immune response (Luplertlop et al., 2011; Sim et al., 2012). Luplertlop et al., 2011 used Digital Gene Expression tag profiling to quantify the impact of DENV infection in SGs. They reported overexpression of Toll pathway components, but did not test their functions.

Instead, they characterized the most abundant protein, an AMP from the Cecropin family, and revealed its broad antiviral function *in vitro*. This Cecropin was not regulated in our transcriptomic analysis, although two other Cecropins (Cecropin A and G) were differentially modulated by the different viruses. Sim et al., 2012 used microarrays to identify SG-specific transcripts as compared to carcasses. An enrichment in immune-related genes suggested the ability to mount an immune response. Further, the same authors determined the transcriptome upon DENV infection. Similar to our data, they reveal a high regulation of serine proteases that could initiate the different immune pathways or play a role in blood feeding when secreted. While they did not functionally test the immune pathways, Sim et al. revealed the antiviral and proviral functions of three DENV-upregulated genes. This supports the complex interaction between gene regulation and function that we also observed. In our study, we functionally tested for the first time the impact of upregulated components from Toll, IMD and JNK pathways on DENV, ZIKV and CHIKV in SGs. JAK/STAT pathway was not activated at the studied time-points. The lack of impact against DENV, ZIKV and CHIKV when silencing the components of Toll and IMD pathways may not reflect the antiviral capabilities of these pathways. Indeed, it is possible that a complete shutdown of the signaling cascades increases virus infection. Importantly, we discovered the antiviral impact of the JNK pathway against DENV, ZIKV and CHIKV in SGs.

JNK pathway activation by DENV, ZIKV and CHIKV in SGs occurred early during infection at 3 dpi. This time corresponds to the onset of SG infection (Salazar et al., 2007). Such an early induction may affect virus dissemination to SGs as reflected in ZIKV and CHIKV infection rates. The JNK pathway can be induced either through IMD-mediated pathogen recognition or oxidative stress. Tak1 activates the IMD and JNK pathways through IKK2 and Hep, respectively (Takatsu et al., 2000). Since Tak1-mediated JNK activation is transient, lasting less than one hour (Park, 2004), the observed JNK activation (lasting 3-7 dpi) is probably due to an IMD-independent activation. Upon oxidative stress, the JNK pathway is induced through p53 upregulation of *Traf4* that then phosphorylates Hep (Lu et al., 2017; Myers et al., 2018; Sax & El-Deiry, 2003). Activated JNK pathway then induces other oxidative stress-associated genes, such as *FoxO* and *κ-B Ras* (AAEL003817) (Essers et al., 2004). In our transcriptomic data, we reported the upregulation of several oxidative stress-associated genes upon infection including *p53*, *Traf4*, *FoxO*, *CYPs* and *κ-B Ras* (Table S1). These results suggest that JNK pathway is activated in SGs as a result of infection-induced oxidative stress.

Using *in vivo* functional studies, we showed that the JNK antiviral function depends on complement system and apoptosis inductions. The complement system is activated when TEPs bind to pathogens and trigger lysis, phagocytosis (Levashina et al., 2001) or even AMP production (Xiao et al., 2014). TEPs such as *TEP15* and *AaMCR* can restrict DENV in mosquitoes (Cheng et al., 2011; Xiao et al., 2014), although *TEP22* does not (Jupatanakul et al., 2017). Apoptosis was previously reported in SGs infected with different flaviviruses and alphaviruses (Day et al., 1966; Girard et al., 2005; Kelly et al., 2012). Apoptosis antiviral function in mosquitoes is supported by two observations. Firstly, an arbovirus with an apoptosis-inducing transgene was selected out during mosquito infection (O’Neill et al., 2015), and secondly, there is an association between the ability to induce apoptosis and colony refractoriness in different virus-mosquito systems (Clem, 2016; Ocampo et al., 2013; Vaidyanathan & Scott, 2006). These indicate that apoptosis can define vector competence, emphasizing its importance as a target to control transmission. In our study, we reported the antiviral functions of a complement system component, TEP20, and an apoptotic component, Dronc, extending our understanding of the two mechanisms in SGs. Based on a lack of synergistic effect when both TEP20 and Dronc were silenced, we propose that complement system and apoptosis function in the same cascade to reduce virus infection. The complement system could mediate apoptotic cell clearance as in mammals (Fishelson, 2001) and in testes of *A. gambiae* (Pompon & Levashina, 2015). Alternatively, apoptosis-triggered nitration of virus surface could be required to direct TEP binding, as in *Anopheles* mosquitoes infected with parasites (Kumar et al., 2010; Oliveira et al., 2012).

A recent study established a negative association between the presence of efficient *Plasmodium*-killing immune response in mosquitoes and epidemics in Africa, confirming the long-suspected impact of mosquito immunity on epidemiology for arthropod-borne diseases (Gildenhard et al., 2019). Upon close inspection of field-derived *A. aegypti* colony transcriptomes (Sim et al., 2013), we found that DENV-refractory colonies expressed a higher level of *Kay,* c-*Jun* and *TEP20.* This suggests that variation in vector competence among these colonies is partially related to JNK pathway. Consistent with our study, this points to a role of JNK pathway in determining the arbovirus transmission dynamics in the field. Because JNK pathway is sensitive to various microbes (Guntermann & Foley, 2011), differential activation by distinct microbes present in natural habitats may also represent a trigger that influences transmission.

Development of transgenic refractory mosquitoes have gained prominence in preventing arboviral transmission (Wilke et al., 2018). Engineered overexpression of a JAK/STAT activator in mosquitoes reduced DENV propagation and established its proof-of-concept (Jupatanakul et al., 2017). However, JAK/STAT is ineffective against ZIKV and CHIKV. To our knowledge, no promising candidates have been identified to be antiviral against DENV, ZIKV and CHIKV. In this context, our work reveals that the JNK pathway components could be harnessed to develop effective transmission blocking tools against a broad range of arboviruses.

## Acknowledgments

We thank the Professor Eng Eong Ooi from Duke-NUS Medical School, for providing the DENV and ZIKV stock, and Professor Lisa Ng from Singapore Immunology Network (SIgN, A*STAR, Singapore), for providing the CHIKV stock. This work was funded by the Tier-3 grant from the Ministry of Education, Singapore (MOE 2015-T3-1-003) and the Duke-NUS Signature Research Programme funded by the Agency for Science, Technology and Research (A*STAR), Singapore, and the Ministry of Health, Singapore.

## Author Contributions

J.F.P. and R.M.K. designed the project. A.C., S.T.T. and B.W. performed the experiments. C.M.M. performed bioinformatics analysis. A.C., C.M.M., D.M., T.V., R.M.K. and J.F.P. analyzed the data. A.C. and C.M.M. made the figures. A.C., C.M.M., R.M.K. and J.F.P. wrote the manuscript. All authors reviewed, critiqued and provided comments on the text.

## Material and Methods

### Mosquitoes

*Aedes aegypti* mosquitoes were collected in Singapore in 2010. Eggs were hatched in MilliQ water and larvae were fed with a mixture of fish food (TetraMin fish flakes), yeast and liver powder (MP Biomedicals). Adults were reared in cages (Bioquip) supplemented with 10% sucrose and water. The insectary was held at 28°C and 50% relative humidity with a 12h:12h light:dark cycle.

### Virus isolates

The dengue virus serotype 2 PVP110 (DENV) was isolated from an EDEN cohort patient in Singapore in 2008 (Christenbury et al., 2010). The Zika virus Paraiba_01/2015 (ZIKV) was isolated from a febrile female in the state of Paraiba, Brazil in 2015 (Tsetsarkin et al., 2016). The chikungunya virus SGP011 (CHIKV) was isolated from a patient at the National University Hospital in Singapore (Her et al., 2010). DENV and ZIKV isolates were propagated in C6/36 and CHIKV in Vero cell line. Virus stocks were titered with BHK-21 cell plaque assay as previously described (Manokaran et al., 2015), aliquoted and stored at −80°C.

### Oral infection

Three-to-five day-old female mosquitoes were starved for 24 h and offered an infectious blood meal containing 40% volume of washed erythrocytes from serum pathogen free (SPF) pig’s blood (PWG Genetics), 5% 10 mM ATP (Thermo Scientific), 5% human serum (Sigma) and 50% virus solution in RPMI media (Gibco). Mosquitoes were left to feed for 1.5 h using Hemotek membrane feeder system (Discovery Workshops) covered with porcine intestine membrane (sausage casing). The virus titers in blood meals were 2 x 10^7^ pfu/ml for DENV, 6 x 10^6^ pfu/ml for ZIKV, and 1.5 x 10^8^ pfu/ml for CHIKV, and validated in plaque assay using BHK-21 cells. Control mosquitoes were fed with the same blood meal composition except for virus. Fully engorged females were selected and kept in a cage with *ad libitum* access to a 10% sucrose solution in an incubation chamber with conditions similar to insect rearing.

### Virus inoculation

Mosquitoes were cold anesthetized and intrathoracically inoculated with either DENV, ZIKV or CHIKV using Nanoject-II (Drummond). The same volume of RPMI media (ThermoFisher Scientific) was injected as control. Functional studies were conducted by inoculating 0.1 pfu of DENV, 0.01 pfu of ZIKV and 0.2 pfu of CHIKV.

### SG collection, library preparation and RNA-sequencing

SGs from DENV- and ZIKV-orally infected mosquitoes were dissected at 14 dpi. Those from CHIKV-orally infected mosquitoes were dissected at 7 dpi. Controls for CHIKV were dissected at 7 days post feeding on a non-infectious blood meal, and at 14 days post feeding for DENV and ZIKV. Inoculum resulted in 100% infected SGs (Supplementary Figure 1). Fifty pairs of SGs per condition were homogenized using a bead mill homogenizer (FastPrep-24, MP Biomedicals). Total RNA was extracted using E.Z.N.A Total RNA kit I (OMEGA Bio-Tek). RNA-seq libraries were prepared using True-Seq Stranded Total RNA with Ribo-Zero Gold kit (Illumina), according to manufacturer’s instructions. Following quantification by RT-qPCR using KAPA Library Quantification Kit (KAPA Biosystems), libraries were pooled in equimolar concentrations for cluster generation on cBOT system (Illumina) and sequenced (150 bp pair-end) on a HiSeq 3000 instrument (Illumina) at the Duke-NUS Genome Biology Facility, according to manufacturer’s protocols. Two repeats per condition were processed. The need for more than two repeats was mitigated by using a large number of tissues from different mosquitoes in each repeat. Raw sequencing reads from RNA-seq libraries are available online under NCBI accessions: SRR8921123-8921132.

### RNA-seq data processing and identification of differentially expressed genes (DEGs)

Reads were quality checked with FastQC (www.bioinformatics.babraham.ac.uk) to confirm that adapter sequences and low-quality reads (Phred+33 score < 20) had been removed. The reads were then aligned against the *A. aegypti* genome [VectorBase(Giraldo-Calderón et al., 2015)] AaegL3.3) using TopHat v2.1.0 (Kim et al., 2013) with parameters –N 6 –read-gap-length 6 –read-edit-dist. 6 set to account for regional genetic variation between the RNA-seq and genome. SAM tools v0.1.19 (Li et al., 2009) and HTSeq v0.6.0 (Anders et al., 2015) were used to format and produce count files for gene expression analysis. DEGs were identified using DESeq2 (Love et al., 2014) with at least 1.4-fold change between control and infected conditions, an adjusted False Discovery Rate (FDR) of 0.05. edgeR (Robinson et al., 2010) and Cuffdiff 2 (Trapnell et al., 2013) (following the Cufflinks pipeline (Trapnell et al., 2012)) were also used to identify DEGs following the same criteria.

### Gene annotations

DEGs were annotated through several pipelines. Predicted gene protein sequences were searched against the NCBI nr database (accessed July 2017) (BLASTp; e-value 1.0E-5 and word size 3), VectorBase (Giraldo-Calderón et al., 2015) *Aedes* peptide database (downloaded July 2017) with BLAST+ (Camacho et al., 2009) (BLASTp; e-value 1.0E-5 threshold and word size 2), and FlyBase (Gramates et al., 2017) *D. melanogaster* peptide database (downloaded August 2017) with BLAST+ (BLASTp; e-value 1.0E-5 threshold and word size 2) for identification of orthologues. Gene ontology terms were assigned in BLAST2GO (Conesa et al., 2005) (program default parameters). Signal peptides were identified using SignalP v4.1 (Petersen et al., 2011). Functional annotations were also assigned based on literature.

### Double-stranded mediated RNAi

Mosquito cDNA was used to amplify dsRNA targets with gene specific primers tagged with T7 promoter as detailed in Supplementary Table 2. The amplified products were *in vitro* transcribed with T7 Scribe kit (Cellscript). dsRNAs were annealed by heating to 95°C and slowly cooling down to 4°C using a thermocycler. Three to five-day-old adult female mosquitoes were cold-anesthetized and intrathoracically injected with 2 or 4 µg of dsRNA using Nanoject II. The same quantity of dsRNA against the bacterial gene *LacZ* was injected as control (dsCtrl). Validation of gene silencing was conducted 4 days post injection by pooling 10 SGs or 5 midguts.

### Gene expression quantification using RT-qPCR

Total RNA was extracted from 10 SGs or 5 midguts using E.Z.N.A. Total RNA kit I, DNAse treated using Turbo DNA-free kit (Thermo Fisher Scientific), and reverse transcribed using iScript cDNA synthesis kit (Biorad). Gene expression was quantified using qPCR with SensiFast Sybr no-rox kit (Bioline) and gene specific primers detailed in Supplementary Table 2, 3 and 4. *Actin* expression was used for normalization. The reactions were performed using the following cycle conditions: an initial 95°C for 10 min, followed by 40 cycles of 95°C for 5 s, 60°C for 20 s and ending with a melting curve analysis. The delta delta method was used to calculate relative fold changes.

### Quantification of virus genome RNA (gRNA) copies using RT-qPCR

Individual pairs of either SGs or midguts were homogenized with a bead Mill Homogenizer in 350 µl of TRK lysis buffer (E.Z. N. A Total RNA kit I). Total RNA was extracted using the RNA extraction kit and reverse-transcribed using iScript cDNA synthesis kit. DENV gRNA copies were quantified by RT-qPCR using i-Taq one step universal probes kit (BioRad) and ZIKV and CHIKV gRNA copies with i-Taq one step universal sybr kit (BioRad) with primers detailed in Supplementary Table 5. Amplification was run on CFX96 Touch Real-Time PCR Detection System (BioRad) with the following thermal profile: 50°C for 10 min, 95 °C for 1 min, followed by 40 cycles at 95 °C for 10 s, 60 °C for 15 s. A melt-curve analysis was added for Sybr qPCR.

Absolute quantification of gRNA was obtained by generating a standard curve for each virus target. Viral cDNA was used to amplify qPCR target using qPCR primers with T7-tagged forward primer. RNA fragments were generated with T7-Scribe kit and RNA concentration calculated by Nanodrop was used to estimate concentration of RNA fragments. Ten-time serial dilutions were quantified with RT-qPCR and used to generate absolute standard equations. Three standard dilutions were used in each subsequent RT-qPCR plate to adjust for inter-plate variation.

### Statistical analyses and software used

One-way ANOVA and post-hoc Dunnett’s test or unpaired T-test were used to test differences in gene expression and log10-transformed gRNA copies. These analyses were done using GraphPad Prism 5. Z-score was used to test differences in infection rate with www.socscistatistics.com/tests/ztest/. Standard error for sample proportion was calculated with www.easycalculation.com.

