## Supplemental information for "JNK pathway restricts DENV, ZIKV and CHIKV infection by activating complement and apoptosis in mosquito salivary glands"

**Supplementary Information**

**Supplementary Text**

**Common DEGs among DENV, ZIKV and CHIKV**

Our comparative high throughput RNA-seq data revealed 19 DEGs common to DENV, ZIKV or CHIKV infection in *A. aegypti* SGs. They included six commonly upregulated, 11 commonly downregulated and two differentially-regulated DEGs. Commonly upregulated genes included the transcription termination factor *Lodestar* (AAEL018118)[1], *Ribunucleoside disphosphate reductase* (AAEL010691) that is involved in DNA synthesis, two immune genes, namely *Juvenile Hormone inducible (JHI,* AAEL000515*)* and *Dicer2 (Dcr2,* AAEL006794*)*, one oxidative stress responsive gene *Cytochrome6Z9* (AAEL009129), and one gene of unknown function (AAEL003732/EAT44971). DEGs that were commonly downregulated included a digestive enzyme, *MaltaseA1* (AAEL009524), three blood-feeding facilitating genes, namely an *apyrase* (AAEL006347), a *34kDa family secreted protein* (AAEL003600) and a *salivary mucin4* (AAEL003100), *Trypsin3A1* (AAEL007818), one stress-responsive gene *Cytochrome9J26* (AAEL014609), one antimicrobial peptide, *Gambicin1* (AAEL004522), a *venom allergen* (AAEL000793) and three genes of unknown functions (AAEL009081, AAEL004899, AAEL010242). The CLIP-domain serine protease *CLIPB37* (AAEL005431) and a fibrinogen-related pathogen pattern recognition protein (AAEL008646) were upregulated by ZIKV and CHIKV, and downregulated by DENV. The largest proportion of DEGs was virus-specific. Although the difference in collection time between the flaviviruses and alphavirus could play a role, these transcriptome signatures reflect virus-specific regulation in SGs.

**DEGs related to blood-feeding**

Arboviruses are transmitted by mosquito saliva secretion during skin probing[2–4]. Alteration of salivary components can alter blood feeding, and thus enhance transmission by: i) increasing probing attempts and probing time[5]; ii) reducing the production of salivary proteins that directly inhibit virus infection[6]; and iii) enhancing amounts of allergens or other inflammatory compounds that increase infection[7]. We noted that DENV, ZIKV and CHIKV modulated 13, 13 and 27 DEGs related to blood feeding, respectively (Supplementary Table 1). Interestingly, four of these DEGs were commonly downregulated by all three viruses. These included an *apyrase* (*ATP diphosphohydrolase*) (AAEL006347), a *salivary mucin* (AAEL003100), a *34 kDa protein* (AAEL003600) and a putative *salivary secreted peptide gene* (AAEL004899). Two other apyrases (AAEL000575 and AAEL006333) were downregulated by DENV and ZIKV, respectively, while nine other mucins were differentially regulated by the three viruses (Supplementary Table 1). Apyrases hydrolyze ATP and ADP to AMP, preventing platelet aggregation during blood-feeding. Apyrase contents in salivary gland extracts from various mosquito species inversely correlated to probing attempts[8]. Mucins may play a role in lubricating the insertion of mouthparts and influence probing. The 34 kDa protein has been shown to enhance DENV infection of keratinocytes by inhibiting the interferon response[9], and the role of the putative salivary secreted peptide, an orthologue of a tick saliva protein[10], is unknown.

Five D7 family genes were downregulated (AAEL006417, AAEL007394, AAEL006423, AAEL006424, AAEL002726) upon ZIKV or CHIKV infection. These highly abundant mosquito saliva proteins[11–13] scavenge biogenic amines[14] and reduce vasoconstriction, plateletaggregating and pain-inducing properties[14–16]. One of these proteins (AAEL006424) inhibits DENV infection in vertebrates[6].

Three odorant binding proteins (OBPs; AAEL018102, AAEL002587 and AAEL000124) were downregulated by DENV or CHIKV, while another one (AAEL005770) was upregulated by ZIKV. DENV-infected SGs showed increased expression of two other OBPs that are involved in probing initiation[5]. OBPs bind to hydrophobic odorant chemicals and carry them to odorant receptors, thereby influencing olfactory and gustatory signaling[17].

**DEGs related to lipid metabolism**

Enveloped viruses such as DENV, ZIKV and CHIKV interact with host lipid membranes for entry, replication, translation, assembly and egress[18,19]. Flavivirus infection alters lipid profile in mosquitoes[20] and is restricted by lipid biogenesis inhibition in mosquito cells[21]. We found 22 DEGs related to lipid metabolism, indicating a strong alteration of the lipid profile in SGs (Supplementary Table 1). Two homologues of *fatty acid synthase* (*FAS*) (AAEL002228 and AAEL001194), which initiate fatty acid biogenesis, were downregulated by ZIKV and CHIKV infection. On the contrary, genes involved downstream of FAS, such as *elongases* (AAEL009574, AAEL013128), *desaturase* (AAEL003645) and *reductase* (AAEL006774), were upregulated by CHIKV. Enzymes involved in synthesis of phospholipids and sphingolipids were also altered. *Glycerol-3-phosphate acyltransferase* (AAEL012743) that initiates phospholipid biogenesis was downregulated by ZIKV. Two *phospholipase A2s* (AAEL001523, AAEL001528) that hydrolyze the 2-acyl ester, and one *phospholipase D* (AAEL000264) that hydrolyses the basic head group were upregulated by CHIKV. *Sphingomyelin synthetases* (AAEL004710, AAEL001381) were upregulated by CHIKV, while *sphingolipid phospholipase* (AAEL003402) was downregulated upon DENV infection. Interestingly, the gene associated with lipid droplets (AAEL005951), induced by DENV infection in mosquito Aag2 cells[22], was downregulated by both DENV and CHIKV in mosquito SGs.

**DEGs related to immune effectors**

AMPs are the hallmark of immune activation and can be regulated independently or in combination by different pathways (Fig. 1a, Supplementary Fig. 6 and 7)[23–25]. Strikingly, *Gambicin1* (*Gam1*) was downregulated by the three viruses (Fig. 1c; Supplementary Table 1), indicating immune inhibition. However, Gam1 shows no effect on DENV replication[26]. Defensins (*DefA* and *DefD*) and cecropins (*CecA*, *CecB* and *CecG*) that have been reported to antagonize arboviruses[27,28] were upregulated upon ZIKV infection. On the contrary, upon DENV infection *CecG* was downregulated. DENV inhibition of *CecG* expression was associated to sfRNA in SGs[29]. It would be interesting to test whether the discrepancy in *CecG* expression between DENV and ZIKV is due to variation in sfRNA function.

The complement system can eliminate pathogens by opsonization, phagocytosis or lysis[30] and is regulated by different immune pathways. Leucine-rich repeat (LRR) proteins guide the binding of TEPs to pathogens and activate the complement system[31]. In *A. aegypti*, TEP15 mediates an anti-DENV response[32]. We observed the upregulation of several TEPs, including *TEP15*, by ZIKV and CHIKV. However, 17 LRRs showed virus-specific differential regulation (Fig. 1 and Supplementary Fig. 7; Supplementary Table 1), suggesting virus-specific variation in complement activation in SGs.

**DEGs related to apoptosis**

In *A. aegypti*, the core apoptotic pathway is activated by Dronc and Ark, which are repressed by Inhibitor of Apoptosis Protein1 (IAP1)[33,34]. Upon activation, Dronc cleaves effector caspases, such as Caspase8, which then degrade physiological substrates, leading to apoptosis[34]. Upstream of the core pathway, IAP1 is inhibited by a set of proteins including Head involution (Hid). *Hid* transcription is induced by p53[35] and Forkhead box O (FoxO)[36], the latter being activated by Protein kinase C53E (Pkc53E)[37,38]. Alternatively, caspase-independent apoptosis is initiated by death executioner bcl2 (Debcl2)[39]. A DENV refractory strain of *A. aegypti* showed higher *Dronc* expression in the midgut, and silencing of the gene significantly increased the susceptibility of this strain to DENV infection[40]. Interestingly, *Dronc* silencing reduced midgut infection of *A. aegypti* when challenged with Sindbis virus (Alphavirus)[41]. This suggests that infection depends on a fine regulation of apoptosis.

We found 27 DEGs related to apoptosis and autophagy in DENV or CHIKV infection (Supplementary Table 1). Some pro-apoptotic factors like *Caspase8* (AAEL014348) and *p53* (AAEL007595) were upregulated, while other pro-apoptotic factors such as *Pkc53E* (AAEL001108), *FoxO* (AAEL012847) and *Debcl2* (AAEL001515) were downregulated by DENV or CHIKV infection. Pro-apoptotic factors, *Ark* (AAEL000874) and *Dronc* (AAEL011562), and anti-apoptotic factor *IAP1* (AAEL009074) were upregulated only by CHIKV. Such contrasting regulatory mechanisms may be triggered due to vector-virus counteractive interactions, where the vector induces apoptosis as a defense against arbovirus, while viruses may block the apoptotic signaling[42]. Autophagy, which is reported to be a pro-viral phenomenon, intertwines with apoptosis[43,44]. Two autophagy related genes (*ATG14*,

AAEL001133; *ATG18A*, AAEL013063) were upregulated upon CHIKV infection. Thus, apoptosis is tightly-regulated upon arbovirus infection in SGs.

**Supplementary Figures**

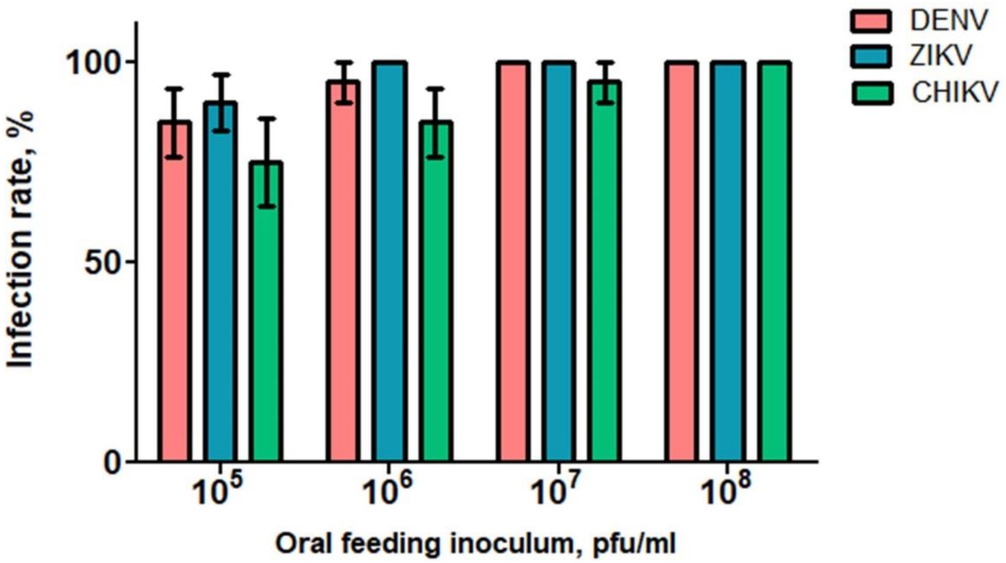


**Supplementary Fig. 1.**

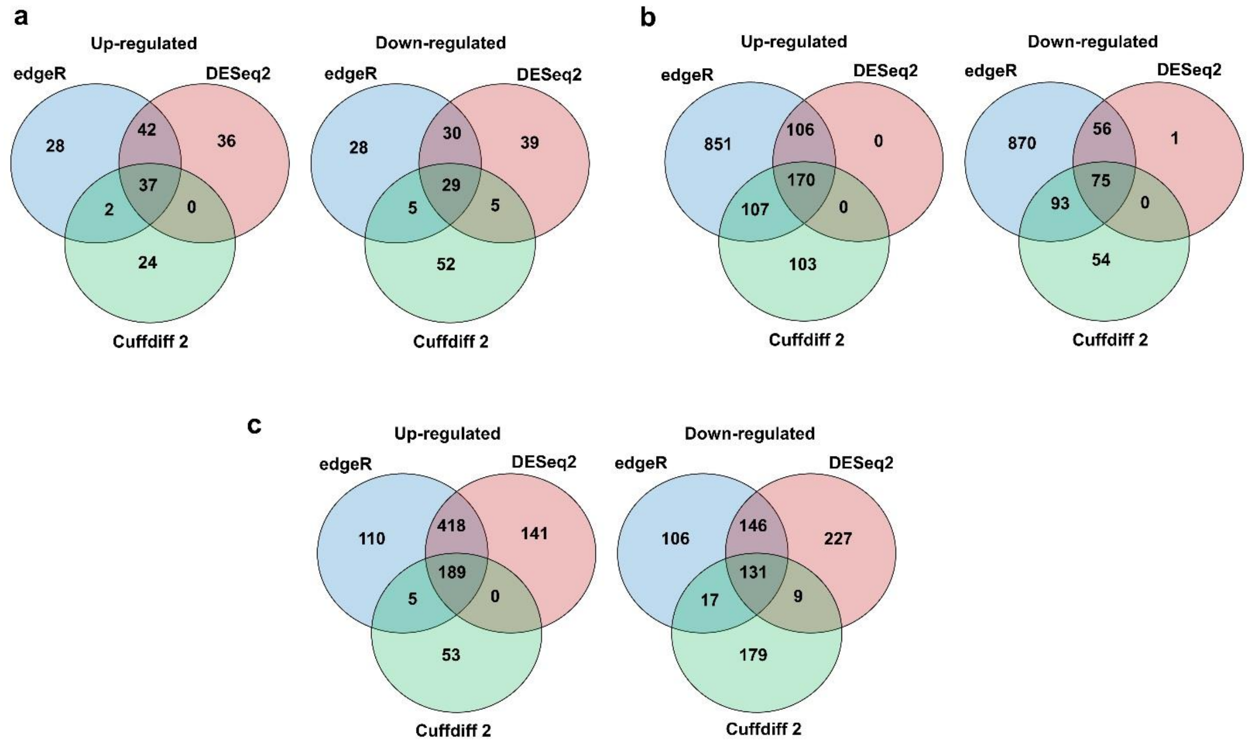


**Supplementary Fig. 2.**

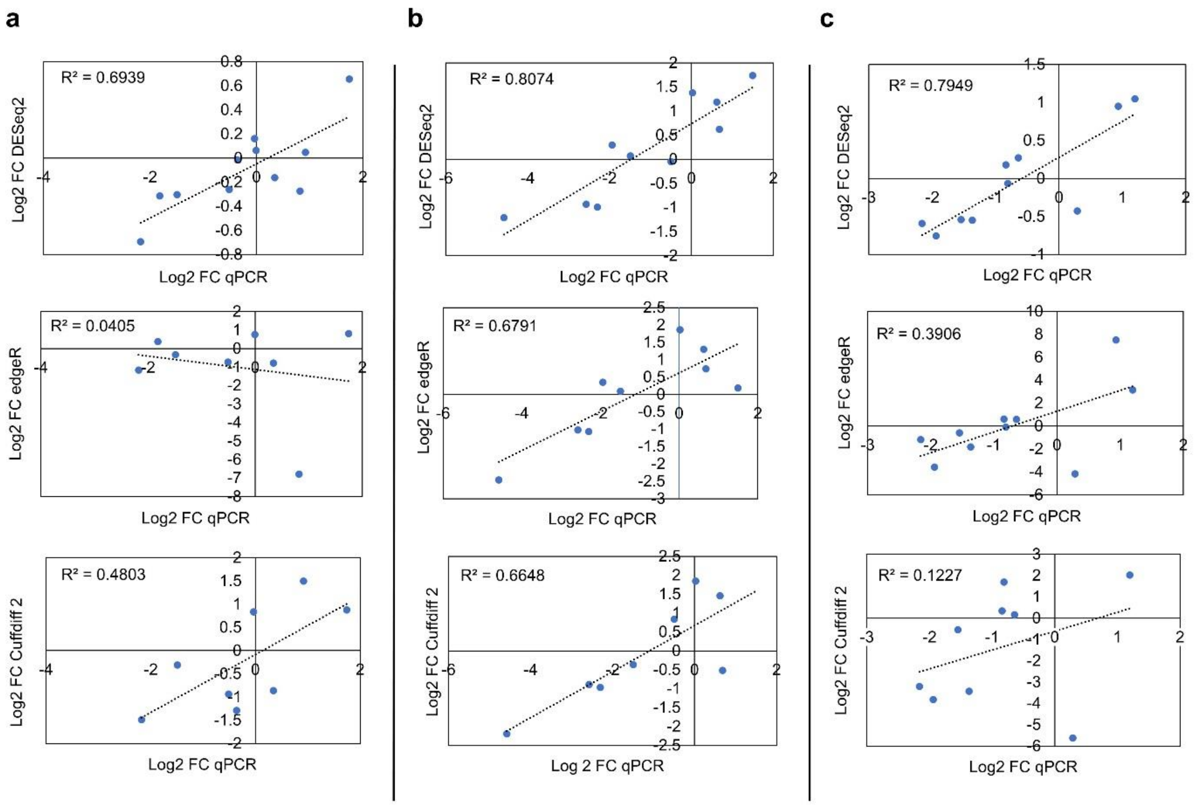


**Supplementary Fig. 3.**

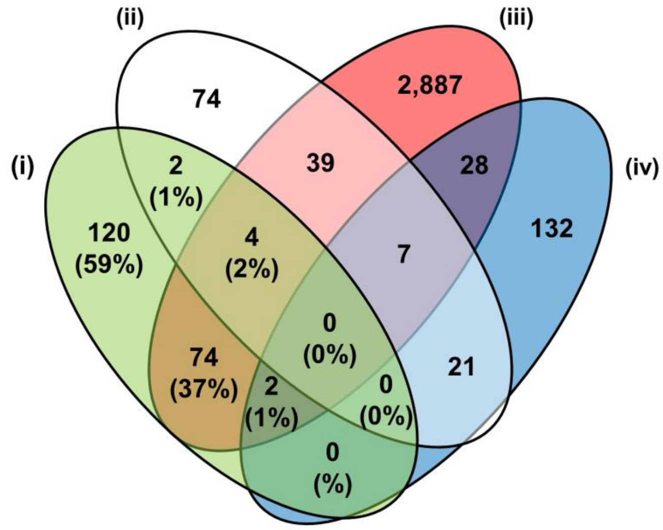


**Supplementary Fig. 4.**

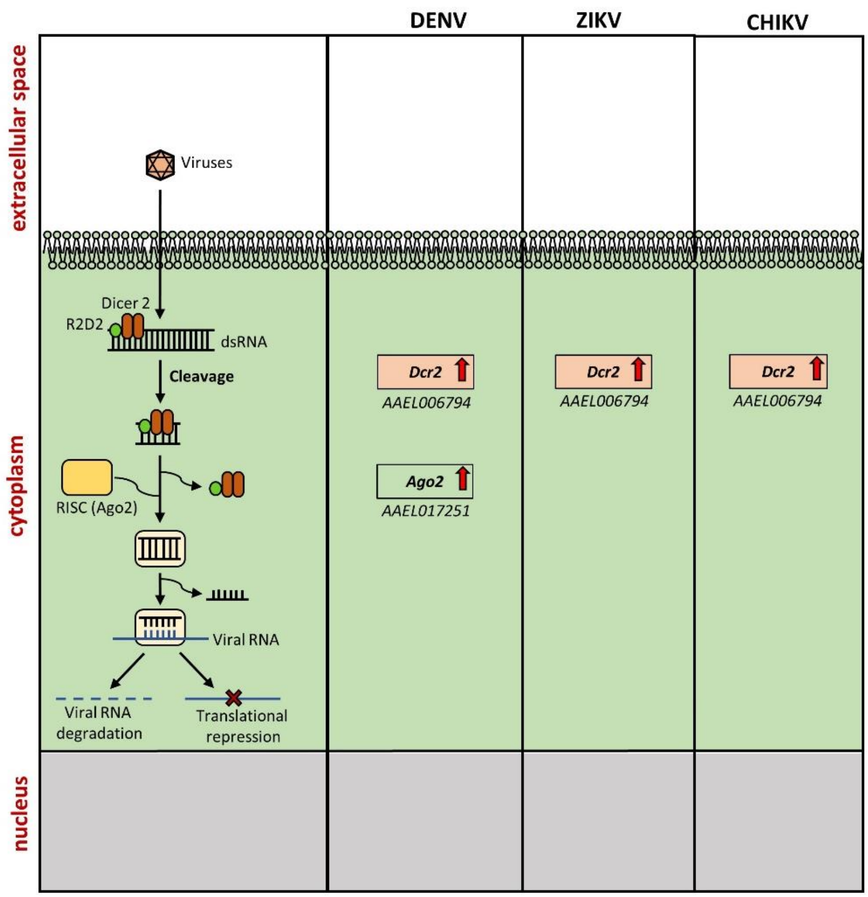


**Supplementary Fig. 5.**

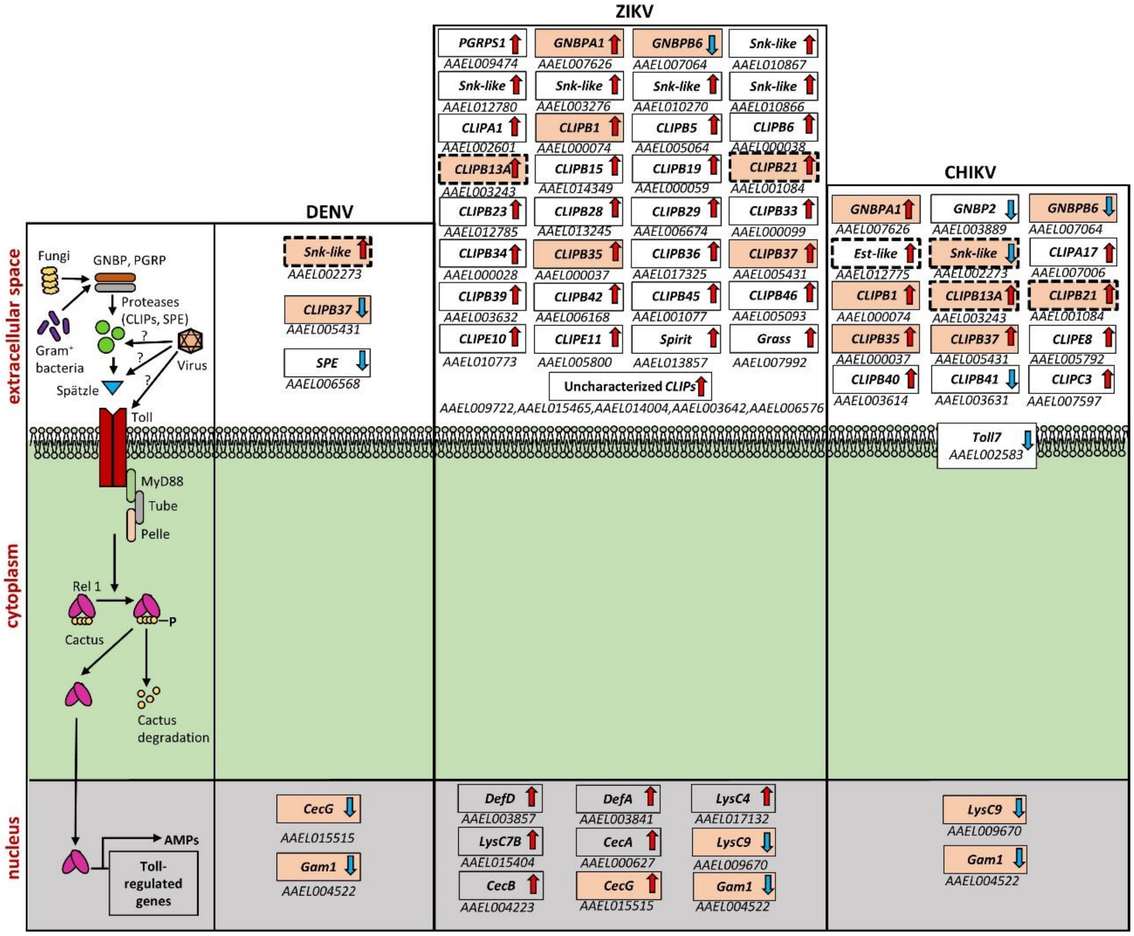


**Supplementary Fig. 6.**

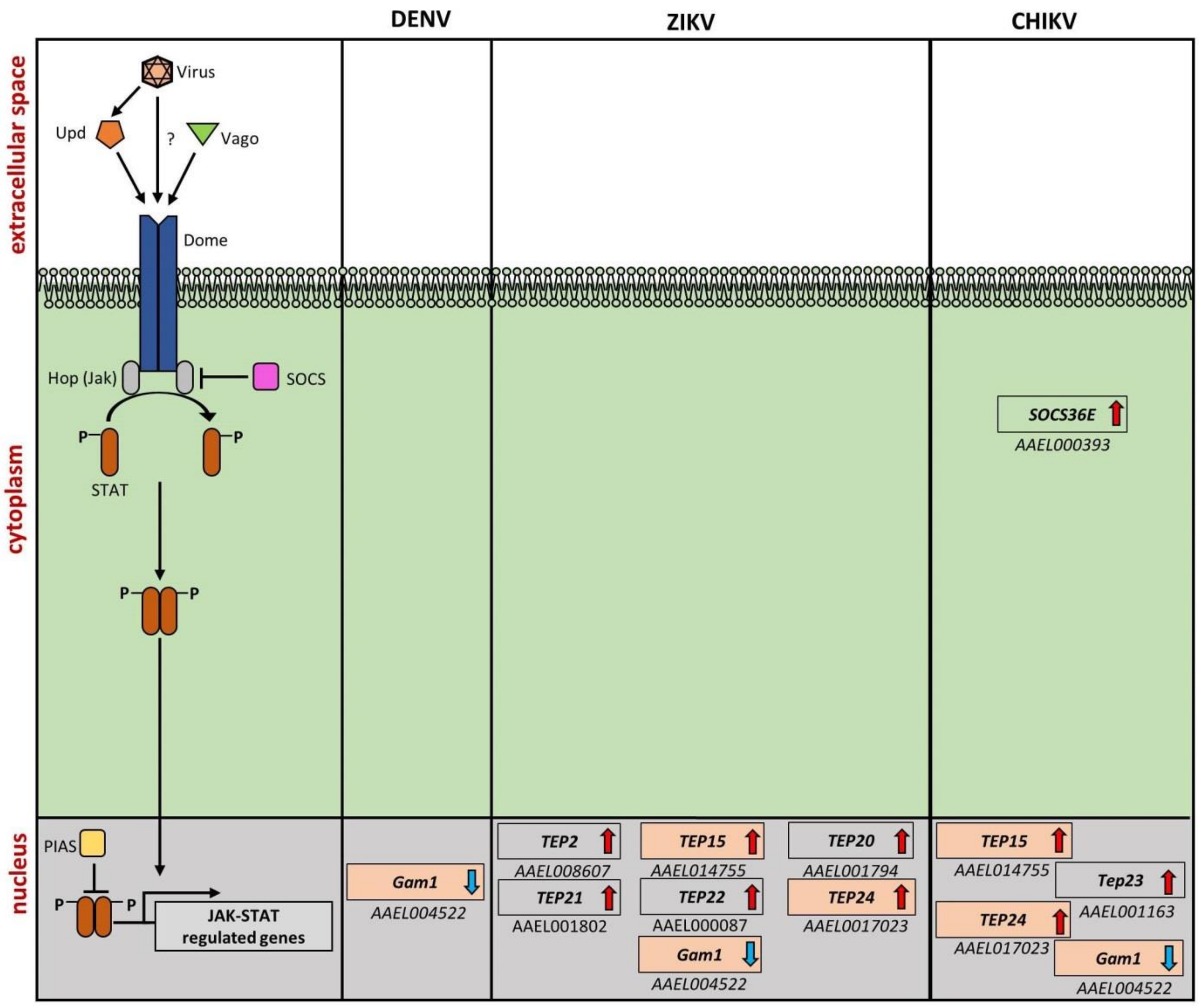


**Supplementary Fig. 7.**

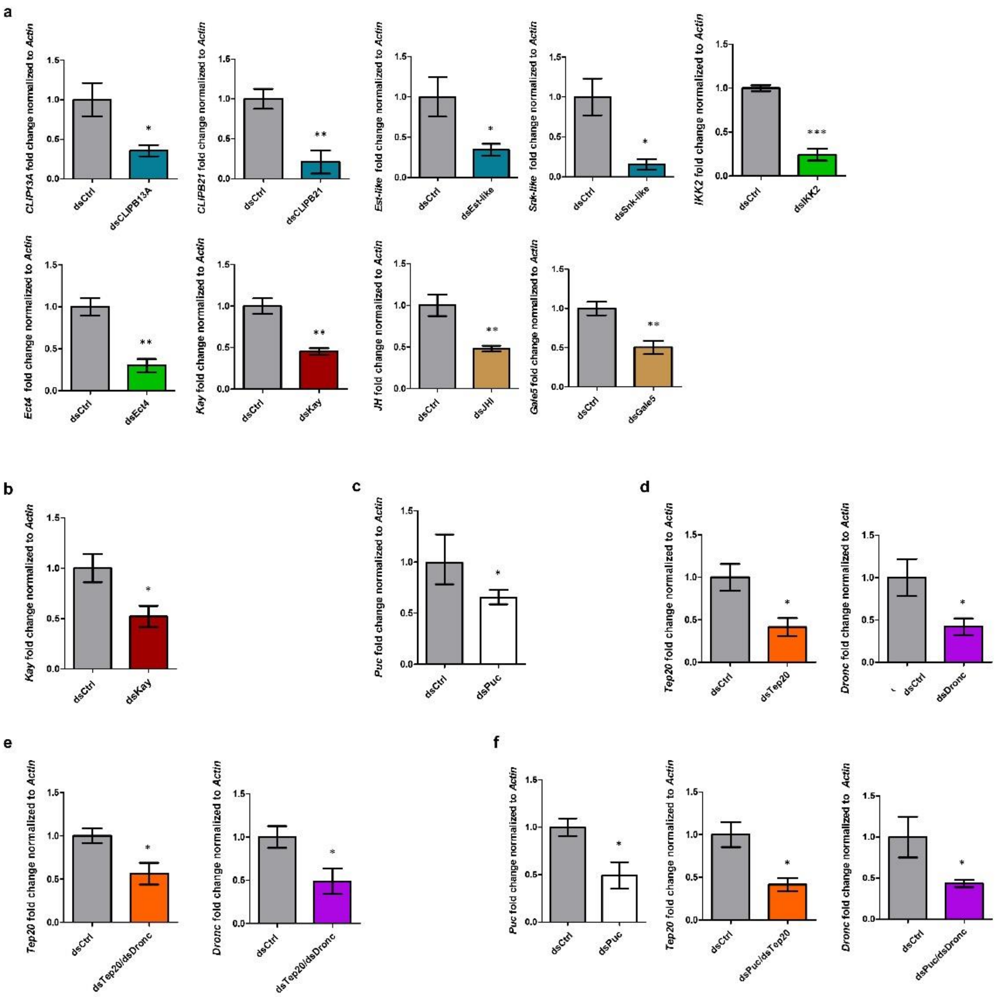


**Supplementary Fig. 8.**

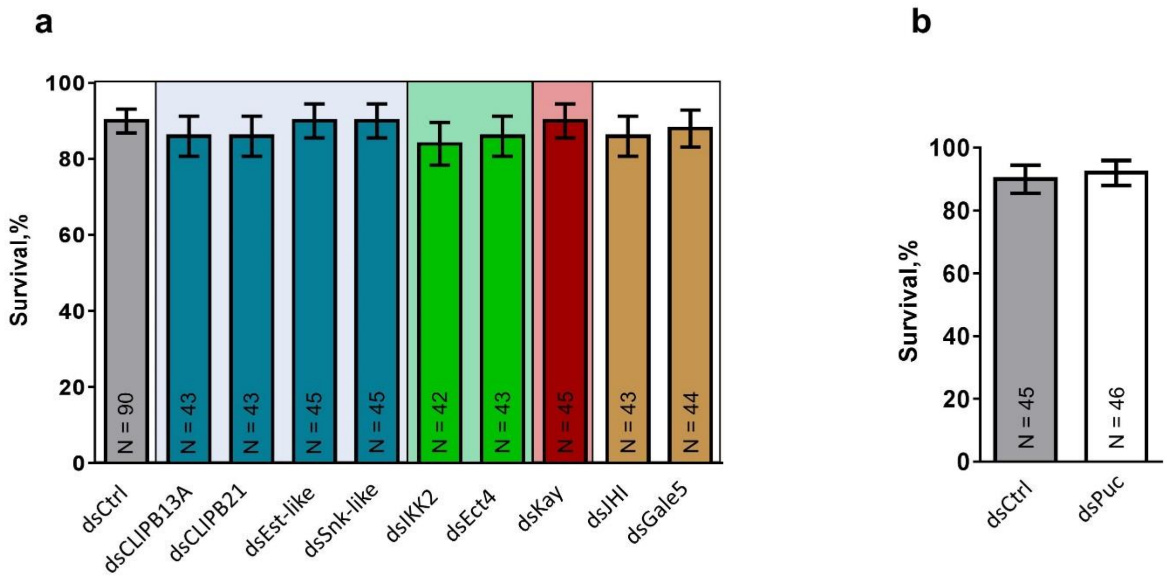


**Supplementary Fig. 9.**

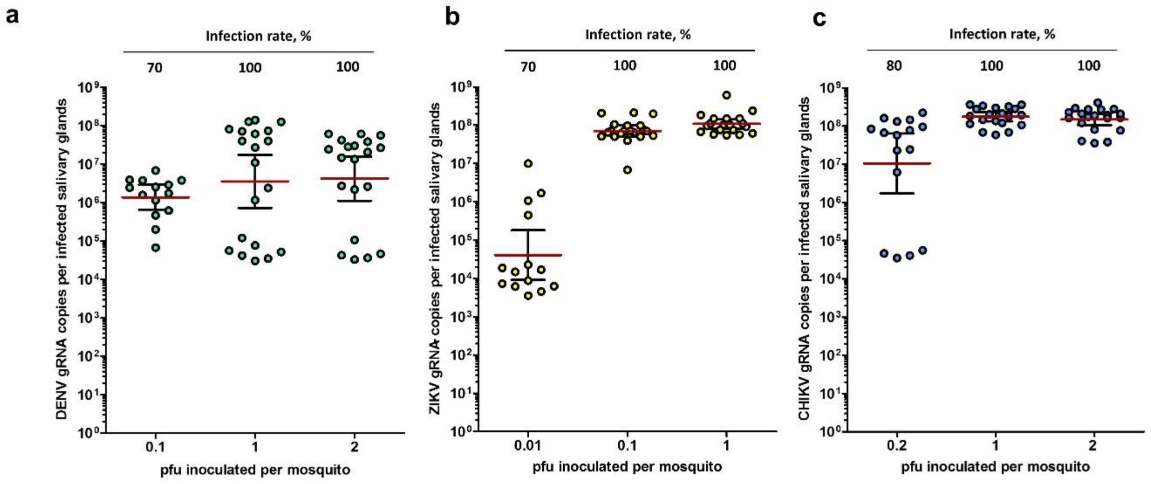


**Supplementary Fig. 10.**

**Supplementary Table 1. Fold changes and functional annotations for DEGs in SGs upon infection with DENV, ZIKV and CHIKV. (Excel sheet)**

**Supplementary Table 2. Primer sequences for candidate gene RNAi silencing and RT-qPCR quantification**

| Gene name/ Acc. No. | dsRNA primers | RT-qPCR primers |
|---|---|---|
| *Actin* / AAEL011197 | | Fw: GAACACCCAGTCCTGCTGACA |
| | | Rv: TGCGTCATCTTCTCACGGTTAG |
| *Lac Z* | Fw: TACCCGTAGGTAGTCACGCA | |
| | Rv: TACGATGCGCCCATCTACAC | |
| *Kayak* / AAEL008953 | Fw: GCCTCCTTTGACGGCTTAC | Fw: CCTCACCGATAGCTTGGACA |
| | Rv: CTGCGCTACGGCTACGTC | Rv: GCGGTAGATTCACACTGGTC |
| *Snake-like* / AAEL002273 | Fw: GTGGTGCGTTTGGGAGAGTA | Fw: CATGTCGATCGTCCACGAAG |
| | Rv: GCACAGATCTGCGAATCAAT | Rv: ACTCACCTTCCAGAGCTTCC |
| *Easter-like* / AAEL012775 | Fw: CGCCATATTCGAGTTCCCT | Fw: AATAGTGCTTGGGTGCTTGC |
| | Rv: CAACCCAATGTCGTTGGAA | Rv: CGGTGTGGTACAGGATGGAT |
| *CLIPB21* / AAEL001084 | Fw: AGGACGGCTCGGAGAAGTA | Fw: GGTGGAGGACTGATGGTTCA |
| | Rv: GATTCCTCGTTCAAACAGGCT | Rv: TTCCAGTAGGCCGTGACATT |
| *CLIPB13A* / AAEL003243 | Fw: CAGTGCGTACTCCGAGGTG | Fw: ATAGAACGAGACTGCGCTGA |
| | Rv: GCAGCCCTTTTGATGTAATTG | Rv: GTCGAACGTTTGCCTCTGAA |
| *IKK2* / AAEL012510 | Fw: AGTGTCATCATGGGCGAAAC | Fw: CGTGGCGAAGAATTTGGAGT |
| | Rv: ACGTTTGTCCTGTTCTGCG | Rv: AGGGTGTTCAGTTCACGGAT |
| *Gale5* / AAEL003844 | Fw: ATCCGCATCAACCAAGCTAC | Fw: GGTGGCTGTGTGATCCATTC |
| | Rv: GCTGCTGTGGAGGGTTGTTA | Rv: GGTACATATGGTGGCGGTTG |

| | | |
|---|---|---|
| *Juvenile Hormone Inducible* / AAEL000515 | Fw: ACCTTGATGGCCGAGAGAAT<br>Rv: CCTGCCATTTTCGTATTTGA | Fw: TTATGAAAGGCGACGAAGCG<br>Rv: TCTCGGACACCAATCTTCCC |
| *Ectoderm Expressed4* / AAEL014931 | Fw: GCAATTGCTCAGGCACAAC<br>Rv: GCCATCGGCATCTTTTCTT | Fw: AAGTGCCACACAAACTGTCC<br>Rv: CCCATTCTCGTACGTCCTCA |

### Supplementary Table 3. Primer sequences for DEG validation

| Virus infection | Gene Accession No. | Primer sequences |
|---|---|---|
| DENV | AAEL017469 | Fw: ACTACTGTGCCGGATTCTGT |
| | | Rv: ATTGGACTGAACAGCAGAGC |
| | AAEL000080 | Fw: TCTCCGACAACTCGGATTTCA |
| | | Rv: TTCTTGCCGAGAAGGGAGTT |
| | AAEL006704 | Fw: CATCTGGGTTGGTACTCCGA |
| | | Rv: TCTCGTCAGAACGATACGTGA |
| | AAEL006953 | Fw: CCTGCTGCTGAAGATCAACC |
| | | Rv: CGGAGTTTCCATTGGAGCTG |
| | AAEL003844 | Fw: CAGGCTATCCCATTGGCAAC |
| | | Rv: GTGGGTAGCTTGGTTGATGC |
| | AAEL012441 | Fw: TGGATGTGGAGATTGCCGAT |
| | | Rv: GGTTGTGCCACACGTTAGTT |
| | AAEL007191 | Fw: GGTGTGATCTTGCCTCTGGA |
| | | Rv: AGCACCATGGCTTTGTTGAG |
| | AAEL003203 | Fw: CGGTTTCGCATTCCTGCTAT |
| | | Rv: AACACGCACAGAATGATCCG |
| | AAEL015121 | Fw: TGCACGCTTTCAGAAACTCC |
| | | Rv: TTTGAGAACTCGCAGTGGGT |
| | AAEL006123 | Fw: ATGGAGCCATCACCATCGAA |
| | | Rv: GAGTGCACAATCGAAGGCAT |

| | | |
|---|---|---|
| | AAEL006498 | Fw: CGCGATGTTTGGATCACTGT |
| | | Rv: CAAGGCACCATTGTTGGTCA |
| | AAEL006259 | Fw: CTCCAACCTGCTAGTGGTCA |
| | | Rv: CCAAGGCAAGCGTAGACTTC |
| ZIKV | AAEL001673 | Fw: CTTCTCTTCCGGTGAAAGGC |
| | | Rv: CCGGATCCGTTATCAACGAC |
| | AAEL002759 | Fw: CTGATGAAGTGTCCCGCAAG |
| | | Rv: ACCGACGACCTTCAACTCTT |
| | AAEL006704 | Fw: CATCTGGGTTGGTACTCCGA |
| | | Rv: TCTCGTCAGAACGATACGTGA |
| | AAEL008283 | Fw: ATGGACGACCTCTCAGTTGG |
| | | Rv: TGGGATGATCACATGCCAGT |
| | AAEL004783 | Fw: TGGTGGTGATGTCCCACAAT |
| | | Rv: TCGGTGTCGTCATAATCGGT |
| | AAEL014937 | Fw: AGCCTTTGCATTTCCACCTG |
| | | Rv: CAGACGACTTGTGGTTAGCG |
| | AAEL004382 | Fw: GCGTTCAAGCTCCGTAATCA |
| | | Rv: CCAGCGATGGGATCAGGAAA |
| | AAEL000886 | Fw: AACTCCTGGCCATCGTAGTC |
| | | Rv: CACCACATTTCGAAGCCAGT |
| | AAEL005676 | Fw: AATTTGATTGACGCCGTCCT |
| | | Rv: GGCAGTCATCCACCAGTTTC |
| | AAEL000647 | Fw: GAGCGAAGAAGAGGTTACGC |
| | | Rv: CACGGCTGAACTTCTAAGCA |
| | AAEL007818 | Fw: AGTTTACCGGCTACCGCATA |
| | | Rv: GCTGAAACTTGGCTGAGTCC |
| CHIKV | AAEL003596 | Fw: TGGACGTGGAAGTTGATGGA |
| | | Rv: CGTGGAAGGTAATCTCGATGC |

| | |
|---|---|
| AAEL005331 | Fw: TAGCGCACTCTTTGTTCACG |
| | Rv: GCGATTCTGCTGGACAAGTT |
| AAEL006050 | Fw: AAGTGGCTGATCCGTTTGTG |
| | Rv: CGTCGTGTCAAAGCCATTCT |
| AAEL003118 | Fw: AAGCCTCACCAGCATCAGAA |
| | Rv: TGTCACTCTTGGCAACAAGC |
| AAEL013341 | Fw: ATATGTTGCCAAGCGGTCAC |
| | Rv: GAAGCGATTTCTTCGGTGCT |
| AAEL006129 | Fw: CCATACGCATCCGTTGATCC |
| | Rv: CGGTTGAACAGCGACTCATT |
| AAEL008646 | Fw: AGTTTAAGGCACGAGGACCA |
| | Rv: CCGAACTTGGTTTGCTCACA |
| AAEL001627 | Fw: TTCCCACTGGATGGAAGCAT |
| | Rv: TAGCAATCGTCGCTGATCCT |
| AAEL001169 | Fw: GCTGACGGATGAAGACATCG |
| | Rv: GCTGACGGATGAAGACATCG |
| AAEL011499 | Fw: GCTTTGATTGACCCACCGTT |
| | Rv: CAGCTTGGTTCCCTCCATTG |
| AAEL002661 | Fw: GGAAGACGGTTCGTGTTCAG |
| | Rv: TCTCTCCGCACATCTCCATC |
| AAEL009750 | Fw: TCTACGACTATGGTGACTGCT |
| | Rv: CGATGCTGGTTCGGAATTGT |
| AAEL003060 | Fw: CCGTCATTCGTGTGGACAAA |
| | Rv: GGAAGCACGACTTTCCTCAG |

**Supplementary Table 4. Primer sequences for JNK pathway-controlled gene expressions and corresponding RNAi silencing**

| Gene name/ Acc. No. | dsRNA primers | RT-qPCR primers sequences |
|---|---|---|

| | | |
|---|---|---|
| *Dronc* / AAEL011562 | Fw: GGGGTGGTCTTCATCGTG | Fw: AGCTGATACGGAGACCGAAG |
| | Rv: GGTCTCCGTATCAGCTTTGG | Rv: GGGTACCGTCGAGAAGCATA |
| *TEP20*/ AAEL001794 | Fw: CGGCATTCGACTGGTTAGC | Fw: CGACCTCTCGCTGTCTACTT |
| | Rv: TTTATCAAGAAACTCGGTGCG | Rv: CAACTGGTTTGCTGTCCCAA |
| *Caspase8* / AAEL014348 | | Fw: CGAATGCCGACTTCCTGTTT |
| | | Rv: GCACAACTCTCGGATGAACC |
| *TEP2* / AAEL008607 | | Fw: CGGCAAGCATCGAGGATAAG |
| | | Rv: ATCAGACTTGCCGAAAGCAC |
| *TEP15*/ AAEL014755 | | Fw: ATCCCAGAAGCCGACAAGAA |
| | | Rv: CGACGACAGTGTCCTGAGTA |
| *TEP24*/ AAEL017023 | | Fw: ATTTCGCTGGTGGAATGACG |
| | | Rv: GGGTCCAAACAAATGTCGCT |
| *ATG14* / AAEL001133 | | Fw: AGTGGTTTCTCGAGCAGTGA |
| | | Rv: TTCCCATTTCCTGGCAAAGC |
| *ATG18A* / AAEL013063 | | Fw: ACGAGGACATACGAATCGCT |
| | | Rv: TTGCAGATCTCCGTCCCTTT |
| *Puc* / AAEL010411 | Fw: TACTTAATCACCCGACCGTTG | Fw: GCACCAGAACATCAAGCAGT |
| | Rv: CGTGTTTGTTGTCACTTGC | Rv: CTGATACCGGCTTGACAGTG |

### Supplementary Table 5. Primer sequences for virus quantification

| **Virus name** | **RT-qPCR primers** |
|---|---|
| CHIKV | Fw: AAAGGGCAAACTCAGCTTCAC |
| | Rv: GCCTGGGCTCATCGTTATTC |

| | |
|---|---|
| DENV | Fw: CAGGTTATGGCACTGTCACGAT |
| | Rv: CCATCTGCAGCAACACCATCTC |
| | Probe:/5HEX/CTCTCCGAGAACAGGCCTCGACTTAAA/3BHQ1/ |
| ZIKV | Fw: CCGCTGCCCAACACAAG |
| | Rv: CCACTAACGTTCTTTTGCAGACAT |
